# Design of a biomolecular adaptive controller to restore sustained periodic behavior

**DOI:** 10.1101/2024.10.11.617962

**Authors:** Frank Britto Bisso, Subham Dey, Guy-Bart Stan, Christian Cuba Samaniego

## Abstract

Periodic behavior is a widespread biological phenomenon occurring across various spatiotemporal scales, where upstream stimuli are encoded into dynamic intracellular signals such as oscillations, varying in duration, amplitude, and frequency. Disruptions to this periodicity can lead to a range of pathologies, for which we propose an adaptive feedback controller with an Incoherent Feedforward Loop (IFFL)-like topology, based on chemical reactions, designed to restore sustained oscillations in systems that have lost their periodicity. By approximating the controller’s dynamics, we defined the design requirements for the first implementation of a biomolecular adaptive controller and tested its applicability for destabilizing the steady–state behavior of a self-inhibiting gene. Numerical simulations illustrate the adaptive behavior of the controller and its ability to tune both the amplitude and the period of the resulting oscillations.

## I. Introduction

Periodic behavior is ubiquitous in biological systems across different spatiotemporal scales, as cellular function often requires regulators to be activated or inhibited repetitively at specific locations, for certain durations, and at defined concentrations [1]. For instance, during neural development, precursor cells encode differentiation messages in the frequency and fold-change of effector (signaling) molecules which exhibit oscillatory dynamics [2]. Similarly, during vertebrate embryogenesis, the sequential formation of the periodic pattern in our spinal cord is considered to be regulated by the interaction between a *segmentation clock* and morphogen gradients [3]. Although the benefit of this kind of dynamic regulation is not fully understood, the absence of periodicity is strongly correlated with various pathologies [4].

In this work, we propose a biomolecular adaptive control strategy (Fig. 1–A) to restore sustained oscillations in a system that has lost its periodicity. Our work is inspired by that proposed in [5], wherein the authors designed an adaptive controller with a similar structure to a delayed feedback controller, but replacing the delay block with a low–pass filter. Building on our previous work, which introduced the design of a band-pass filter using an Incoherent Feedforward Loop (IFFL) network [6], we now propose the design of a biomolecular adaptive controller with a similar topology.

**Fig. 1.**
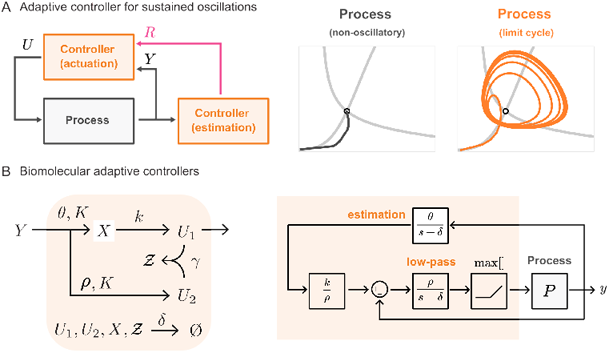
Adaptive feedback controller. The left side of Panel A shows a closed–loop diagram with an adaptive controller decoupled in two blocks: actuation and estimation. In this work, we took as a control design problem the generation of sustained oscillations, illustrated in the right side of Panel A. The left side of Panel B presents the schematics of the chemical species that comprise the adaptive controller. The chemical reactions involving those species, following Eq. (2), are mapped to the component of the block diagram presented at the right of Panel B through the help of Eq. (7).

The rest of the manuscript is structured as follows: Section II–A introduces the self–inhibiting gene as our study case. Sections II–B and C detail the controller’s network and examine its behavior in isolation. In Section II–D, the closedloop system at steady state is analyzed, considering the self–inhibiting gene as the process. Section II–E demonstrates the controller’s capability to generate sustained oscillations and its adaptive behavior in the presence of parametric perturbations. Conditions for oscillatory behavior are established in Section II–F through a local stability analysis. Ultimately, the adaptive controller’s ability to tune the period and amplitude of the resulting oscillations is numerically characterized.

## II. Results

Throughout this paper, we indicate chemical species with capital letters (e.g *U*) and their corresponding concentration with the corresponding lowercase letter (e.g *u*).

### A. Problem formulation

In this work, we address *sustained oscillations*, defined as those occurring when the dominant eigenvalues of the Jacobian matrix, evaluated at equilibrium, are complex-conjugates with a positive real part. A fundamental requirement for this kind of periodic behavior is the presence of negative feedback [7]. Hence, the simplest biological oscillator will be a self–inhibiting gene with an appropriate delay, introduced through a series of downstream processes [8]. By neglecting this delay, we can simulate the disruption of an oscillator into a non–oscillatory behavior.

This network considers a single gene that produces a repressor *Y* that downregulates its own production with a rate constant *αp*, and decays at rate δ, and can be modeled with the following chemical reaction:

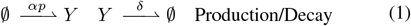

where 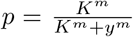. Thus, with the adaptive controller, we intend to destabilize its steady–state behavior into a limit cycle.

### B. Adaptive feedback control formulation

Fig. 1–A introduces a general diagram of a closed–loop system for the adaptive controller. In this setup, we emphasize the absence of a user defined reference for tracking, as the controller is able to estimate the reference from the dynamics of the process. For visual purposes, we decoupled the adaptive controller into two components: an “estimation” module, and an “actuation” module which compares the process output with the estimated reference and actuates over the process accordingly.

As for the design, described in Fig. 1–B, an IFFL motif takes an input *Y* and downregulates the production of species *U*_2_ and *X* with rate constants *ρp* and *θp*, respectively. Then, the *X* species produces *U*_1_ at rate constant *k*. Likewise, *U*_2_ downregulates the expression of *U*_1_ through sequestration at a rate defined by *γ* via an intermediatory complex called *Z*. Ultimately, we assumed that every species decays at rate δ. We can describe this network through the following chemical reactions:

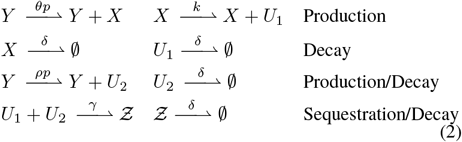

Following the law of mass action, we can model these chemical reactions, and can also have an equivalent non–dimensionalized representation. We define the new variables, 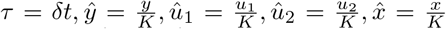 and parameters 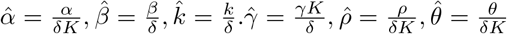, resulting in the following Ordinary Differential Equations (ODEs):

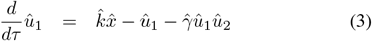

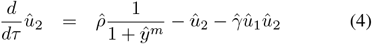

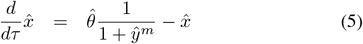

We can also rewrite the dynamics of the self–inhibiting gene driven by the controller actuation with a non–dimensionalized representation,

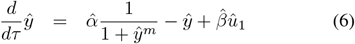

### C. Approximation of the controller dynamics

In this subsection, we find an approximate expression of the dynamics of the controller in isolation when the sequestration regime is fast, defined as 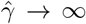.Now, we introduce the following transformations: 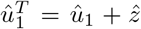 and 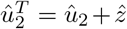, which results in the following set of equations: 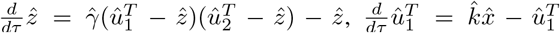 and 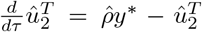, where *y*^*^ represents the input to the controller in isolation. Following a similar analysis given in [6], which have its roots in the principles of time scale separation, we can write 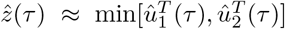 and 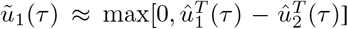 in the fast sequestration region.

#### Theorem 1.

*The expression of the controller’s dynamics in the fast sequestration regime, when working in isolation involving Eqs*. (3)*–*(5), *and with the help of the suitable transformation stated in the aforementioned subsection, can be written as*

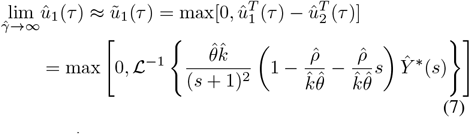

*where ŷ* ^*^(*s*) *represents the Laplace transform of the input to the controller in isolation, and* ℒ^−1^ *represents the inverse transform. Furthermore*,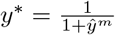.

*Proof*. The Laplace transform of 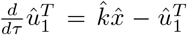 yields 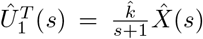,where 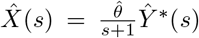.From Eq. (5), we obtain 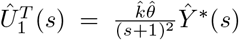 with *ŷ* ^*^(*s*) as the Laplace transform of *y*^*^. Whereas, the Laplace transform of 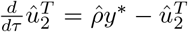 can be written as 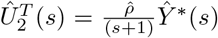. Thus, subtracting 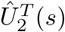 from 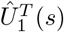 and taking the inverse Laplace transform results in the expression of the approximated controller dynamics when working in isolation at the fast sequestration regime.

In Fig. 2, we compare the approximated dynamics of *û*_1_(*τ*) from Eq. (7) (black dashed lines) with the dynamics obtained from solving Eq. (3)-(5) for increasing sequestration rates, and considering a unit step and a sinusoidal input, both with zero initial conditions. The provided simulations show that, in a fast sequestration regime, we can legitimately approximate the dynamics of *û*_1_(*τ*) using the expression in Eq. (7). Previous works in the literature have shown that adaptive control techniques like time delayed control can be used for the stabilization of an unstable system [5]. However, we are aware that time delayed control leads to an infinite dimensional system, so an alternative approach is to use a low pass filter and derivative controller. Accordingly, we define 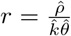 as the adaptive metric. Then, by setting *r* = 1, we can interpret Eq. (7) as the computation of a low-pass filter and a temporal derivative. As a remark, the transfer function derived in Theorem 1 doesn’t describe the full dynamics of the system, but was rather used to approximate *û*_1_(*τ*) and show the presence of the adaptive metric in the expression of controller’s dynamics.

**Fig. 2.**
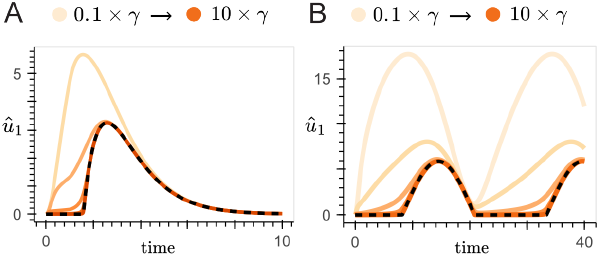
Approximated dynamics. We compared the prediction from Eq. (7), in blacked dashed line, for different sequestration rates for a (A) unit step and (B) sinusoidal input. The non–dimensionalized ODEs (Eq. 3–5) were used for these simulations, and the corresponding nominal parameters were 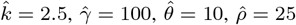 and 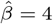.

We can further map the kinetics of the chemical reactions in the fast sequestration regime into components of the block diagram detailed in Fig. 1–B, through the help of Eq. (7). The “estimation” module is equivalent to the top block corresponding to a proportional gain *θ* with a low–pass filter with cutoff frequency δ. This estimated reference (*r*^*^) is scaled by a factor of *k/ρ*, and compared with the current state of the process, generating an error 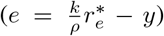.

After a low–pass filter (*e*^*^), the error passes through a non–linear operation max[0, *e*^*^], taking only the positive values of this error. From Eq. (7), the Laplace transform variable *s*, which represents the derivative function in the Laplace domain, has a negative sign. Then, *û*_1_ can only respond to the positive part of that function, which occurs when the gradient is negative and the derivative term in Eq. (7) becomes positive. Conversely, for positive gradients, this part becomes negative, and if it dominates the sign, the output of the controller becomes zero.

### D. Boundedness and Steady-state analysis

In this subsection, we proof that there is a unique steadystate value for the system dynamics given by Eq. (3)–(6), and that the trajectories described by those equations are bounded. Following the same approach as [6], we can write 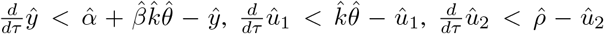 and 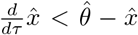 which ensures boundedness of the non–dimensionalized state variables *ŷ, û*_1_, *û*_2_ and 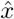, respectively.

Furthermore, we can find the nullclines for the controller by setting Eqs. (3)–(5) to zero. This leads to

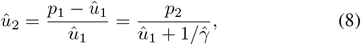

where 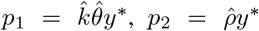,and *y*^*^ = (1 + *ŷ*^*m*^)^−1^. The values *ū*_1_ and *ū*_2_ are steady state values of the non–dimensional model parameters *û*_1_ and *û*_2_ respectively. From, Eq. (8), we can construct a second order polynomial in the variable *û*_1_, given by 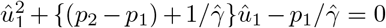.We can rewrite 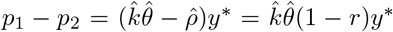 with the help of adaptive metric mentioned in Subsection II–B 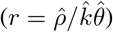. Then, the only valid solution of the equation of the second order polynomial is given by,

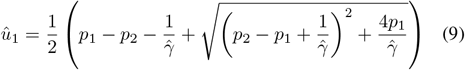

Analyzing its derivative with respect to *ŷ* we obtain,

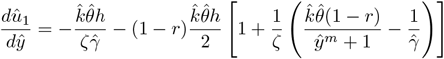

where *h* = *mŷ*^*m*−1^*/*(1 + *ŷ*^*m*^)^2^ and 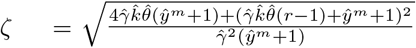, which is ζ *>* 0, ∀*r*.This function is monotonically decreasing 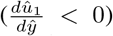 for *r <* 1 and approaches zero for high values of 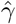 when *r* = 1. Similarly, by making Eq. (6) equal to zero, we can find the process’ nullcline as

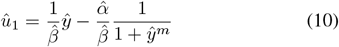

Analyzing its derivative with respect to the input 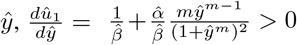. Therefore, this function is monotonically increasing 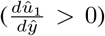. Then, for zero initial conditions and *r* ≤ 1, both Eq. (8) and Eq. (10) intersect at a single equilibrium point.

### E. Analysis of sustained oscillations

The schematics of the interaction between the self-inhibiting motif (process) and the adaptive controller are depicted in Fig. 3–A. The generation of sustained oscillations from the non–oscillatory dynamics of the process can be observed in Fig. 3–B, in orange and black, respectively, numerically validating our design. Moreover, Fig. 3–C illustrates the phase plane between the species *ŷ* and *û*_1_. The two grey lines correspond to the nullclines of the system, computed using Eq. (8) and (10). Both intersect in a single point, around which the limit cycle is generated.

**Fig. 3.**
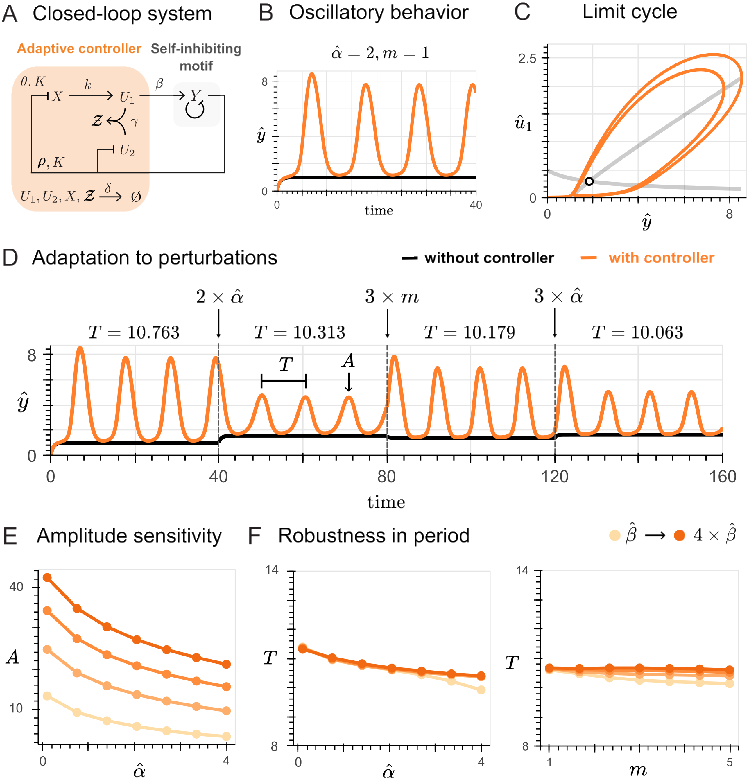
Analysis of sustained oscillations. Panel A shows the closedloop diagram for our study case. Panel B demonstrates the generation of oscillations through the application of the controller. Panel C shows the nullclines of the closed–loop system and a limit cycle. Panel D shows the adaptation of the closed-loop system when the process undergoes disturbances such as an production 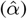 and a increased cooperativity (*m*). Panel E and F systematically characterize how the amplitude and period of the resulting oscillations change with respect to the process, for increasing values of actuation gain 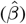. The variable *A* was determined as the mean amplitude of the resulting oscillations, and the variable *T* was estimated as the mean time difference between adjacent peaks, both calculated over a simulation time of *τ* = 200. The non–dimensionalized ODEs (Eq. 3–5) were used for these simulations, with the same nominal parameters as Fig. 2.

Furthermore, to address the adaptive behavior of the proposed feedback controller, we introduced perturbations to the process by simulating the closed-loop system under different values of production rate 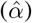 and cooperativity (*m*). Fig. 3–D illustrates how the controller is able to reject disturbances and maintain sustained oscillations. Extending this result, for different fixed values of gain that guarantee sustained oscillations, Fig. 3–E shows the decay of the amplitude, suggesting sensitivity in this feature. Nevertheless, within the same conditions, Fig. 3–F characterizes the relative robustness in the period. For instance, by setting the gain to twice its nominal value, a maximum of 10% change in period (from 11.4 to 10.4) is expected, upon a four-fold increase of the process’s production rate 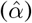.

### F. Deriving the oscillatory condition

#### 1) Linear stability analysis

To evaluate the stability of the closed–loop system, we take a cue from the Routh–Hurwitz stability criteria. We can write down the Jacobian for the non–dimensional model as

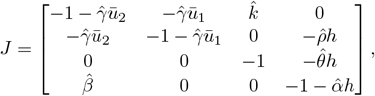

Where 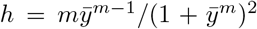, and 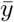 is the steady state value of the variable *ŷ*. The expression of the characteristic polynomial can be obtained by calculating the determinant of the matrix *P*_*a*_(*s*) = (*sI* −*J*), where *I* is an identity matrix of appropriate dimension.

##### Theorem 2.

*The conditions that ensure sustained oscillations in the fast sequestration regime for the linearized system dynamics, as the closed loop system have bounded trajectories, are given by*

- *Condition a:*

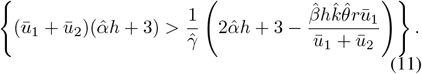
- *Condition b:*

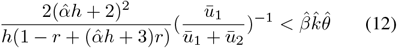

*Proof*. We calculate the determinant of matrix *P*_*a*_(*s*) and obtain the characteristic polynomial 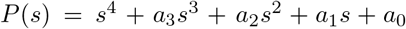, where 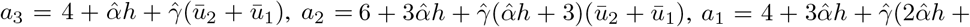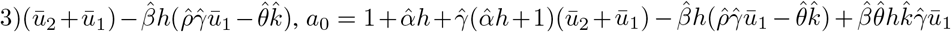. The total number of roots of the characteristic polynomial is 4. In order for sustained oscillations to exist, we need to have complex conjugate pair of roots with positive real part, since a negative real part leads to damping oscillations. We then construct the Routh— Hurwitz table, and find some sufficient conditions for the system to admit two unstable eigenvalues (a complex pair), considering the fast sequestration regime.

**TABLE 1.** Routh—Hurwitz Table.

|  |  |  |  |
| --- | --- | --- | --- |
| $s^4$ | 1 | $a_2$ | $a_0$ |
| $s^3$ | $a_3$ | $a_1$ | 0 |
| $s^2$ | $b_1$ | $a_0$ | 0 |
| $s^1$ | $c_1$ | 0 | 0 |
| $s^0$ | $a_0$ | 0 | 0 |

Following the Routh–Hurwitz stability criteria, to determine the oscillatory condition, we can analyze the signs of *b*_1_ and *c*_1_ when 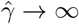.The signs of *a*_3_ is always positive due to design constraints, while the sign of *a*_0_ was found to be always positive in the fast sequestration regime and desired adaptive metric as crosschecked from numerical simulations. Then, any change of sign in the first column of the Routh table will be determined by *b*_1_ and *c*_1_, which are defined as 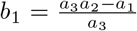, and 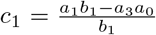. Particularly, in order to have two sign changes in the the first column of the Routh table, we need to have *b*_1_ as positive and *c*_1_ as negative, as crosschecked from numerical simulations. This leads to 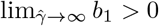. Hence, it must be that 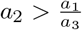. In the fast sequestration regime, 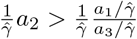, where

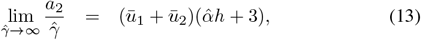

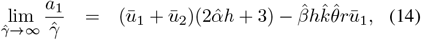

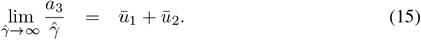

Thus, 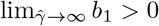 when

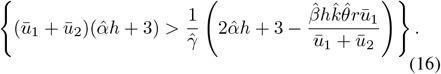

Following similar steps, in the fast sequestration regime, we can find *c*_1_ *<* 0 when 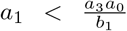 or 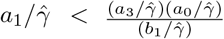. Thus,

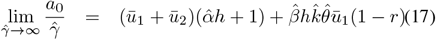

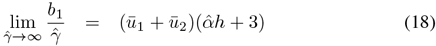

Thus, 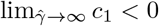 when

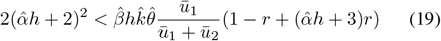

Hence, from Eqs. (16) and (19), we can say the theorem is proved.

In Fig. 4, we fixed the adaptive metric and screened over different values of 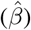 with respect to increasing values of the production rate 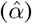. The plane was then divided between 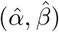 pairs such that when evaluating the Jacobian at their corresponding equilibrium point, it results in at least one pair of complex–conjugate eigenvalues with a positive real part (orange) and those which either present a negative real part or no complex eigenvalues at all (grey). We further crosschecked with simulations that the grey area is only composed of sets of eigenvalues with one pair of complex–conjugates with a negative real part. The black dashed line represents the oscillatory condition derived from Eqs. (11) and (12), which matches the inferior color boundary of the numerical simulations for high sequestration rates, with increasing accuracy as the adaptive metric approaches one (see Fig. 4–B2). The light orange region represents where the divergence of the nonlinear system from the linear system analysis proposed in this subsection can be visualized. The local linear analysis provides eigenvalues with only a real part, suggesting conditions where no oscillations occur in the system trajectories. However, numerical testing of the nonlinear system behavior in these regions reveals sustained oscillations.

**Fig. 4.**
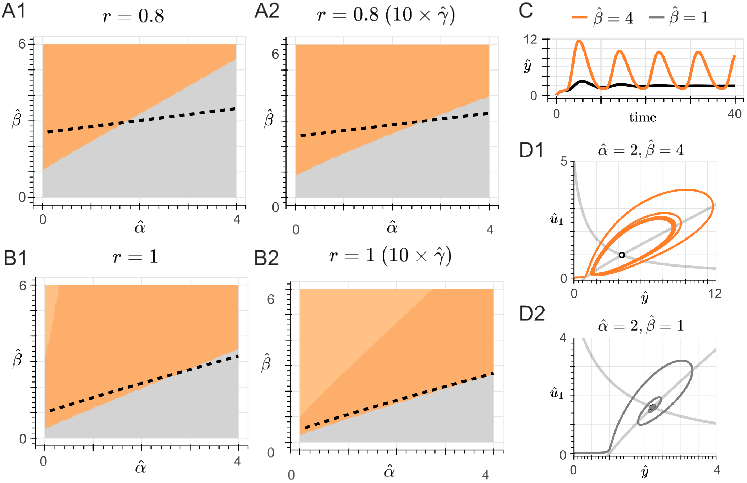
Oscillatory condition. We evaluated the existence of complex eigenvalues with a positive real part for different values of and 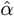, colored 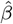 in orange. Eigenvalues with all negative roots or complex–conjugates with a negative real part were colored in grey. The black dashed line defines the oscillatory condition derived from Theorem 2. Panels A1 and B1 show the results for increasing the values of adaptive metric, while Panels A2, and B2 illustrate the convergence of the simulations with the oscillatory condition for a higher sequestration rate. Panel C exemplifies the dynamics of the closed–loop system when the gain 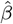 do (orange) or do not (grey) satisfy the oscillatory condition. Panel D1 and D2 show the limit cycle and stable fixed point for that same comparison, respectively. The non–dimensionalized ODEs (Eq. 3–5) were used for these simulations. The same nominal parameters as Fig. 2 were used, except for 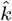, which was defined as 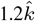 for Panels A1, A2, C, D1 and D2.

Moreover, to gain further insights from Theorem 2, based on Eq. (9), we can determine the fraction between the sequestration species *û*_1_ and *û*_2_ at steady–state for 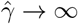, leading to

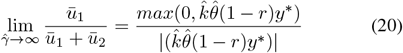

From this expression, we notice that the adaptive metric can admit values so that *r* ≤ 1. In this range, given that Condition a) holds true as crosscheked from numerical simulations, we notice that Condition b) from Theorem 2 simplifies to an effective gain condition 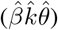 which, if satisfied, guarantees the oscillatory behavior. For instance, Fig. 4–C compares the dynamics of the closed–loop system for the nominal process parameters and two different gain values, above and below the threshold defined by Eqs. (11) and (12). In grey, a decaying oscillation is shown, characterized for its convergence to a stable fixed point in the phase plane (Fig. 4–D2). Likewise, in orange, a sustained oscillation is obtained, as well as its corresponding limit cycle (Fig. 4–D1). On the other hand, from Eq. (20), we notice that, for values of *r >* 1, the inequality in Eq. (12) would not be satisfied, resulting in the absence of oscillations.

#### 2) Sensitivity analysis

We acknowledge the limitations for experimentally finetuning the rate 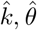 and 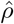, so that *r* is exactly 1. Eq. (20) suggests that the adaptive metric can be relaxed, as long as *r* ≤ 1 provided that 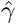 is sufficiently high. Thus, in Fig. 4–A1 and A2, we show that relaxing the adaptive metric up to 20% of its ideal value result in the oscillatory behavior for an appropriate value of gain 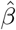, although the oscillatory condition derived from the local stability analysis becomes less precise. Within that error margin, we numerically evaluated the response of the system to variations in the controller’s parameters (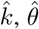and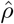). The gain was set as the minimum value obtained from the oscillatory condition. In Fig. 5–A, we show how those variations affect the amplitude and period of the oscillations. Overall, lower values of *r* are correlated with higher amplitude, shorter periods and the addition of a phase.

**Fig. 5.**
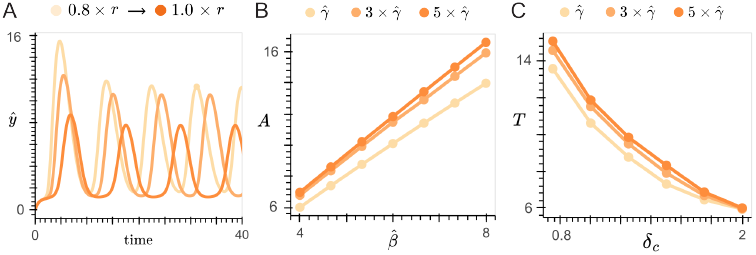
Modulating amplitude and period. Panel A shows the effect of varying the controller’s parameters 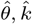 and 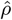 in the amplitude and period of the resulting oscillations within a 20% deviation from the ideal adaptive metric. Panel B characterize the tunability of the amplitude through the gain for different fixed values of sequestration rate, while Panel C does through different degradation rates. The non–dimensionalized ODEs (Eq. 3–5) were used for these simulations, with the same nominal parameters as Fig. 2, except for Panel C, where the ODEs without the variable change detailed in Section II–B were used. The nominal parameters for this case were *k* = *ρ* = 2.5, *γ* = 1000, *θ* = δ = 1, *K* = 0.1, and *β* = 4. The amplitude (*A*) and period (*T*) of oscillations were calculated with the same method as Fig. 3.

To further expand on the possibility of independently controlling the amplitude and period of the resulting oscillations, we increased the gain value and measure the amplitude of oscillations for different fixed sequestration rates. This resulted in a proportional relationship between gain and amplitude, as shown in Fig. 5–B. Moreover, considering the original (dimensionalized) ODEs, we defined a specific degradation rate constant (δ_*c*_) for the controller’s species. For Fig. 5–C, we increase the degradation and measure the period of the resulting oscillations for different, fixed sequestration rates. In this case, an inversely proportional relationship was found between the specific degradation and the period.

## III. Discussion

To the best of our knowledge, this work introduces the first implementation of an adaptive control strategy based on chemical reactions. Our findings on the design of synthetic gene networks used for controlling systems whose dynamics are partially understood or unknown - a common situation in biology - contributes with a new operation to the repertoire of IFFL functions, which include fold-change detectors [9], pulse generators [10], gradient sensors [11], compensators [12], molecular band-pass filters [6] and feedback controllers [13]. Our design represents the simplest adaptive controller, which is a derivative that, instead of fine tuning the genetic circuit’s design to the oscillatory region in the parameter space, destabilizes the process [5]. From the approximated dynamics of the controller, derived in isolation and under the assumption of a high sequestration regime, we note that the proposed network resembles asymptotically a Proportional–Derivative (PD) controller. By constraining the rate constants 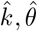 and 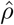, so that 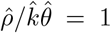, we construct the adaptive controller, following the electronic design proposed in [5]. For generating sustained oscillations using the proposed control strategy, we state Theorem 2, and thus define a design requirement based on the effective gain of the controller 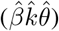.Numerical simulations are provided throughout this work for complementing the theoretical analysis, illustrating the adaptive behavior of the controller when perturbations are introduced and the validity of Theorem 2.

The circadian clock is the only biological oscillator for which the pathological loss of periodicity has been addressed in vitro with optogenetic techniques [14], whereas a pharmacological approach has been demonstrated theoretically [15]. In this case, rather than directly setting gene expression levels, our work illustrates the design of genetic circuits based on a specific dynamic property. Accordingly, for an appropriate gain and specific degradation, the control strategy enables amplitude and period modulation. Synthetic oscillators could also benefit from this architecture, as it allows adaptation to disturbances, compared with current approaches that involve redesigning oscillatory networks for increased robustness [16], [17]. Nevertheless, our design is mainly constraint by the adaptive metric, which could be challenging to meet in a practical setting, although an error margin (*r <* 1) was proof to be admissible. For future work, we plan to extend our analysis of the disturbance rejection feature to account for biological noise, uncertainty, and retroactivity phenomena. Additionally, we aim to explore amplitude and frequency modulation in more complex chemical networks.

## References

[1] J. H. Levine, Y. Lin, and M. B. Elowitz, “Functional roles of pulsing in genetic circuits,” Science, vol. 342, no. 6163, p. 1193–1200, Dec. 2013. [Online]. Available: 10.1126/science.1239999

[2] P. Li and M. B. Elowitz, “Communication codes in developmental signaling pathways,” Development, vol. 146, no. 12, Jun. 2019. [Online]. Available: 10.1242/dev.170977

[3] P. François and V. Mochulska, “Waves, patterns, bifurcations: A tutorial review on the vertebrate segmentation clock,” Physics Reports, vol. 1080, p. 1–104, Aug. 2024. [Online]. Available: 10.1016/j.physrep.2024.05.002

[4] A. Goldbeter, “Dissipative structures and biological rhythms,” Chaos: An Interdisciplinary Journal of Nonlinear Science, vol. 27, no. 10, Sep. 2017. [Online]. Available: 10.1063/1.4990783

[5] K. Pyragas, V. Pyragas, I. Z. Kiss, and J. L. Hudson, “Adaptive control of unknown unstable steady states of dynamical systems,” Physical Review E, vol. 70, no. 2, Aug. 2004. [Online]. Available: 10.1103/PhysRevE.70.026215

[6] Y. Zhang, C. C. Samaniego, K. Carleton, Y. Qian, G. Giordano, and E. Franco, “Building molecular band-pass filters via molecular sequestration,” in 2022 IEEE 61st Conference on Decision and Control (CDC). IEEE, Dec. 2022. [Online]. Available: 10.1109/CDC51059.2022.9993401

[7] B. Novák and J. J. Tyson, “Design principles of biochemical oscillators,” Nature Reviews Molecular Cell Biology, vol. 9, no. 12, p. 981–991, Oct. 2008. [Online]. Available: 10.1038/nrm2530

[8] H. Hirata, S. Yoshiura, T. Ohtsuka, Y. Bessho, T. Harada, K. Yoshikawa, and R. Kageyama, “Oscillatory expression of the bhlh factor hes1 regulated by a negative feedback loop,” Science, vol. 298, no. 5594, p. 840–843, Oct. 2002. [Online]. Available: 10.1126/science.1074560

[9] L. Goentoro, O. Shoval, M. W. Kirschner, and U. Alon, “The incoherent feedforward loop can provide fold-change detection in gene regulation,” Molecular Cell, vol. 36, no. 5, p. 894–899, Dec. 2009. [Online]. Available: 10.1016/j.molcel.2009.11.018

[10] E. Nakamura, C. Cuba Samaniego, F. Blanchini, G. Giordano, and E. Franco, “Design of a sequestration-based network with tunable pulsing dynamics,” Mar. 2024. [Online]. Available: 10.1101/2024.03.24.586474

[11] C. C. Samaniego, J. Kim, and E. Franco, “Sequestration and delays enable the synthesis of a molecular derivative operator,” in 2020 59th IEEE Conference on Decision and Control (CDC). IEEE, 2020, pp. 5106–5112.

[12] R. Du, M. J. Flynn, M. Honsa, R. Jungmann, and M. B. Elowitz, “mirna circuit modules for precise, tunable control of gene expression,” Mar. 2024. [Online]. Available: 10.1101/2024.03.12.583048

[13] M. Whitby, L. Cardelli, M. Kwiatkowska, L. Laurenti, M. Tribastone, and M. Tschaikowski, “PID control of biochemical reaction networks,” IEEE Transactions on Automatic Control, 2021.

[14] M. Foo, O. E. Akman, and D. G. Bates, “Restoring circadian gene profiles in clock networks using synthetic feedback control,” npj Systems Biology and Applications, vol. 8, no. 1, Feb. 2022. [Online]. Available: 10.1038/s41540-022-00216-x

[15] J. H. Abel, A. Chakrabarty, E. B. Klerman, and F. J. Doyle, “Pharmaceutical-based entrainment of circadian phase via nonlinear model predictive control,” Automatica, vol. 100, p. 336–348, Feb. 2019. [Online]. Available: 10.1016/j.automatica.2018.11.012

[16] F. Zhang, Y. Sun, Y. Zhang, W. Shen, S. Wang, Q. Ouyang, and C. Luo, “Independent control of amplitude and period in a synthetic oscillator circuit with modified repressilator,” Communications Biology, vol. 5, no. 1, Jan. 2022. [Online]. Available: 10.1038/s42003-021-02987-1

[17] M. Tomazou, M. Barahona, K. M. Polizzi, and G.-B. Stan, “Computational re-design of synthetic genetic oscillators for independent amplitude and frequency modulation,” Cell Systems, vol. 6, no. 4, pp. 508–520.e5, Apr. 2018. [Online]. Available: 10.1016/j.cels.2018.03.013

